# Whole blood proteome dynamics defines predictive diagnostic and prognostic signatures of cryptococcal infection

**DOI:** 10.1101/2025.05.29.656814

**Authors:** Michael Woods, Jason A. McAlister, Lauren Segeren, Mayara Silva, Jared Deyarmin, Amirmansoor Hakimi, Daniel Hermanson, Jana Richter, Stephanie N. Samra, Jennifer Geddes-McAlister

## Abstract

Across the globe, fungi are impacting the lives of millions of people through the development of infections ranging from superficial to systemic with limited treatment options. To effectively combat fungal disease, rapid and reliable diagnostic methods are required, including current methodologies using antigen detection, culturing, microscopy, and molecular tools. However, the flexibility of these platforms to diagnose infection using non-invasive methods and predict the outcome of disease are limited. In this study, we apply state-of-the-art mass spectrometry-based proteomics to perform dual perspective (i.e., host and pathogen) profiling of cryptococcal infection. Whole blood collected over a temporal scale following murine model challenged with the human fungal pathogen, *Cryptococcus neoformans*, detected >3,000 host proteins and 160 fungal proteins. From the host perspective, temporal regulation of known immune-associated proteins, including eosinophil peroxidase and lipocalin-2, along with suppression of lipoproteins, demonstrated infection- and time-dependent host remodeling. Conversely, from the pathogen perspective, known and putative virulence-associated proteins were detected, including proteins associated with fungal extracellular vesicles and host immune modulation. We also observed and validated a new mechanism of immune system response to *C. neoformans* through modulation of haptoglobin. Further, we assessed the predictive power of dual perspective proteome profiling toward prognostics of cryptococcal infection and report a previously undisclosed integration among virulence factor production, immune system modulation, and individual model survival. Together, our findings pose novel biomarkers of cryptococcal infection from whole blood and highlight the potential of personal proteome profiles to determine the prognosis of cryptococcal infection, a new parameter in fungal disease management.

## Introduction

Fungal diseases impact millions of people across the globe, ranging from superficial infections to systemic diseases with mortality rates exceeding 90%^1–3^. The surveillance of fungal pathogens and infection parameters are challenged by limited methodology, expanding geographical and host niches, and the emergence of new fungal species^4,5^. Traditional diagnostic approaches applied to fungal infections are labor-intensive and time-consuming, requiring microbial culturing combined with identification techniques, such as microscopy^6,7^. Other approaches, such as histopathology and serology, are highly invasive, requiring visual detection and staining of fungal cells from cerebral spinal fluid of a patient. Notably, advances in molecular diagnostics, such as PCR-based detection, sample sequencing, and protein fingerprinting (e.g., matrix-assisted laser desorption/ionization mass spectrometry) have advanced, along with antigen detection of fungal proteins shed within host fluids, provide reliable and robust detection strategies^8^. However, these approaches provide limited information about the state of infection and host status and are typically performed upon suspicion of the final stages of cryptococcal infection (i.e., cryptococcal meningitis) with poor prognostic outcome^9^. Alternatively, high resolution mass spectrometry-based approaches show promise toward diagnosis of cryptococcal infections; however, detection of fungal proteins within the highly complex host matrix demands high specificity and sensitivity, and the promise of diagnostics from non-invasive sources shows potential, but is yet to be fully realized^10,11^.

To capture the importance of these challenges, the World Health Organization developed a fungal pathogen priority list, identifying *Cryptococcus neoformans* as a critical pathogen based on rising rates of infections, disease severity, and increasing occurrence of antifungal resistance^12,13^. This list brought new awareness toward the global impact of fungal disease and justification for renewed interest and commitment to combating fungal infections^14^. For *C. neoformans*, infection is commonly correlated with immune status of the host, with immunocompromised individuals displaying increased susceptibility to infection and elevated mortality^15–17^. For *C. neoformans*, which resides within the natural environment, including decaying tree bark, soil, and pigeon guano, initiation of infection upon inhalation by the host followed by dissemination in the absence of an effective immune response, such as macrophage recognition and fungal cell clearance and T-cell activation, promotes infection^18^. Infection rates within immunocompromised individuals exceeds 220,000 occurrences each year with over 80% of individuals succumbing to infection, and approximately 20% of deaths accounted for in individuals with HIV/AIDS^3,19^. Methods to prevent infection are limited, and while increased availability of anti-viral therapies have reduced the global burden of cryptococcosis, such treatment show increased mortality dependent upon timing of initiation treatment^20,21^. Moreover, strategies to disarm the pathogen by impairing virulence factor production show promise but tend to display reduced efficacy within an immunocompromised host^22–27^. Another hurdle for fungal disease management is a lack of safe and effective therapeutics and dwindling treatment options associated with rising rates of antifungal resistance^28^. Specifically, evolutionary similarity and homology between human and fungal genomes limit the number of druggable targets and lend to increase cytotoxicity within the host. Further, the emergence of fungal pathogens with intrinsic resistance to current antifungals and an increasing prevalence of resistance within the clinic present additional challenegs^29–32^.

Mass spectrometry-based proteomics presents a powerful and adaptable approach to detecting changes in protein production and abundance within complex biological matrices for the diagnosis of disease^33–35^. Moreover, applications toward diagnostics, prognostics, and drug discovery and development are growing^36–38^. Recent expansion of proteomics applications toward infectious disease research and specifically, fungal diseases, has initiated a new avenue of biological discovery and opportunity for improved diagnostic potential^39–41^. Moreover, the integration of advanced bioinformatics platforms and artificial intelligence is expanding the applications of mass spectrometry-based proteomics of infectious disease within clinical research settings^42^. Another emerging area of study within infectious disease research using proteomics is the assessment of infection from dual perspectives (i.e., host and pathogen) to better understand the mechanisms of immune system recognition and activation, along with maintenance of host homeostasis throughout the duration of disease^43,44^. Further, new biological insights into pathogenesis and virulence, identification of putative novel druggable targets, and monitoring pathogen adaptation within the host environment provide new strategies to overcome disease^11,45^. Despite these applications and scientific advances, accurate proteomics-based diagnostic and prognostic markers of cryptococcal infection are still lacking.

In this study, we leverage the power of the novel Thermo Scientific™ Orbitrap™ Astral™ Zoom mass spectrometer with advanced computational pipelines to delve into the dual perspective response of cryptococcal infection. The Orbitrap™ Astral™ Zoom mass spectrometer provides increased ion utilization, scan speed, and quantitative dynamic range for narrow-window data-independent (nDIA) acquisition methodologies, which were originally established using the Orbitrap™ Astral™ mass spectrometer for biological samples, including cell lines and biofluids^46–48^. Using a murine model of cryptococcosis, we extract whole blood over the duration of disease (i.e., 1 day point inoculation [dpi] to experimental end point) to define dual perspective proteome remodeling. We perform nDIA combined with library-based and library-free spectral searches to comprehensively identify changes in production and abundance of host and fungal proteins across temporal and infection scales. We detected over 3,000 host proteins and 160 fungal proteins to define core and modulated dual perspective proteomes from whole blood. From the host perspective, temporal regulation of known immune-associated proteins, including eosinophil peroxidase and lipocalin-2, along with suppression of lipoproteins, demonstrated infection- and time-dependent host remodeling. We also observed and validated a new mechanism of immune system response to *C. neoformans* through modulation of haptoglobin. These data support the discovery of new immune-associated signatures of cryptococcal infection with diagnostic potential to determine the state and severity of disease and efficacy of the host immune response. Conversely, from the pathogen perspective, known and putative virulence-associated proteins were detected, including proteins associated with fungal extracellular vesicles and host immune modulation. Moreover, distinct patterns of fungal protein production proposed new strategies to detect fungal infection as early as 1 dpi, and to monitor the progression of disease across time. Finally, correlation amongst parameters of host and fungal protein abundance integrated with survival patterns, supports a direct connection between diagnostics based on proteome remodeling and prognostics for informing disease management strategies.

## Materials and Methods

### Fungal strains, growth conditions, and media

*Cryptococcus neoformans* var. *grubii* wild-type (WT) strain H99 (serotype A) was maintained on yeast peptone dextrose (YPD) agar plates (2% dextrose, 2% peptone, 1% yeast extract, 1% agar) at 30 °C. For murine assays, *C. neoformans* H99 was grown overnight at 37 °C in YPD, sub-cultured at 1:100 in YPD for approximately 16 h at 37 °C, and collected and washed twice in phosphate buffered saline (PBS). Cells were enumerated with a hemocytometer and resuspended to 4.0 x 10^6^ cells/mL in PBS.

### Murine infection model

Murine infection assays were performed under the approved University of Guelph Animal Utilization Protocol (4193 and 5308), in accordance with all animal handling guidelines. Twenty female BALB/c elite mice aged 6-to-8 weeks (Charels River Laboratories, ON, Canada) were inoculated intranasally with 50 µL of 2 x 10^5^ cells of *C. neoformans* or PBS as a control under isoflurane anesthesia as previously described^49^. The mice were monitored daily for signs of morbidity and euthanized by isoflurane and CO_2_ inhalation upon reaching predetermined endpoints (i.e., >20% body weight loss, respiratory issues, or visible neurological deficits). To adhere to animal handling guidelines (i.e., 50 µL blood collection once per week) and provide a time course of infection, the mice were divided into four groups of five mice each. Saphenous vein blood collection of infected group 1 and uninfected group 1 was collected on days 1, 8, and 15 and saphenous vein blood collection of infected group 2 and uninfected group 2 was collected on days 4 and 11. At the terminal point (i.e., approximately 18 dpi) cardiac blood was collected from all mice immediately after euthanization. Notably, at each time point, a matched uninfected mouse was culled alongside the infected mouse. Samples were flash frozen in liquid nitrogen and stored at -80 °C until processed for proteomic analysis.

### Protein extraction

Blood samples for proteomics were prepared as previously described with minor modifications^50,51^. Briefly, 10 µl whole blood was diluted to 300 µL in 100 mM Tris-HCl (pH 8.5), containing a protease inhibitor cocktail (Sigma-Aldrich), and sodium dodecyl sulphate (2%, final concentration). Samples were probe sonicated in an ice bath (5 cycles, 30 s on/30 s off, 30% power), reduced with 10 mM dithiothreitol for 10 min at 95 °C with shaking at 800 followed by alkylation with 55 mM iodoacetamide for 20 min at room temperature in the dark. Following acetone precipitation overnight at -20 °C, samples were washed and resuspended in 8 M urea/40 mM HEPES for protein quantification by BSA (bovine serum albumin) tryptophan assay. Samples were diluted in 50 mM ammonium bicarbonate, normalized to 25 μg, and digested with trypsin/LysC (Promega, protein/enzyme ratio, 25:1). Digestion performed overnight at room temperature, stopped by addition of 10% v/v trifluoroacetic acid (TFA), and peptides purified by STop And Go Extraction (STAGE) tips^52^. Peptides were dried in a speed vac at 30 °C for 45 min under vacuum conditions until completely dry. Preparation of the cryptococcal samples was performed as outlined above following collection of cell pellets cultured in YPD for 16 h.

Dried peptide samples were resuspended in 2% acetonitrile (ACN)/0.1% formic acid (FA) to an estimated concentration. Peptide concentrations were quantified using the Pierce^TM^ fluorometric peptide quantification assay (Thermo Fisher Scientific) according to the manufacturer’s protocol. Samples were then normalized to a final concentration of 35 ng/µL to ensure equal peptide input for downstream analysis. Indexed retention time (iRT) synthetic peptides (Biognosys AG) were prepared according to the manufacturer’s instructions, diluted 20- fold in 2% ACN/0.1% FA, and spiked into each sample at a fixed volumetric ratio of 1:5 (iRT:sample) prior to LC-MS/MS analysis.

### Mass spectrometry analysis

Whole blood and cardiac blood sample peptides were separated using the Thermo Scientific™ Vanquish™ Neo™ ultra high-performance liquid chromatography system in a trap-and-elute injection configuration and analyzed with the Orbitrap™ Astral™ Zoom mass spectrometer. A Thermo Scientific™ EASY-Spray™ HPLC ES7550PN column (75 μm x 50 cm, 2 μm pore size) was used for analytical separation. Five hundred nanograms of peptides from individual whole blood samples were injected and separated using a 60 samples per day (SPD) chromatographic method with a 23.35-min gradient. The mobile phases were 0.1% formic acid in water (A) and 0.1% formic acid in 80% acetonitrile (B). Details of the liquid chromatography gradient and liquid chromatography configuration are provided (Supplemental Table 1). Eluted peptides were analyzed using nDIA on the Orbitrap™ Astral™ Zoom mass spectrometer. Parameters for full scan MS1 and nDIA MS2 are provided (Supplemental Table 2-4). Six independent gas-phase fractionation injections were performed using pooled samples from whole blood–specific or cardiac blood–infected and uninfected groups, as well as a *C. neoformans* reference culture. All injections used the same sample load and LC gradient for a comprehensive spectral library assembly. Parameters for gas phase fractionation full scan MS1 and nDIA MS2 are provided (Supplemental Table 5-6).

### Mass spectrometry data processing

All mass spectrometer output files were analyzed using Spectronaut v19 (Biognosys AG). A comprehensive spectral library was generated using protein FASTA files against *C. neoformans* var. *grubii* serotype A (strain H99/ATCC 208821; UniProt UP000010091; 7,429 sequences) and *Mus musculus* (17,836 Swiss-Prot reviewed entries, UniProt)^53^. The spectral library was assembled by combining Pulsar search results from individual blood samples, blood gas-phase fractionation runs, and *C. neoformans* gas-phase fractionation runs. This combined library was then used to search the experimental data files. Default search parameters were applied during the DIA analysis search. More specifically, enzyme and modification parameters were set at trypsin enzyme specificity with a maximum of two missed cleavages, carbamidomethylation of cysteines (fixed modification), and oxidation of methionine and N-acetylation of proteins (as variable modifications). Spectral matching of the peptides was performed with a false discovery rate (FDR) of 1% identified proteins with a minimum of two peptides for protein identification. Quantification was based on MS2 peak areas, no imputation strategy was used, and local normalization was applied.

### Bioinformatics

Data analysis and visualization were performed with Perseus (version 1.6.2.2)^54^ and ProteoPlotter^55^. Data was filtered to remove contaminants, reverse peptides, and peptides only identified by site, proteins identified with <2 peptides, and intensities were log_2_ transformed. Valid value filtering with proteins identified in at least 70% of replicates in each group (core proteome) or at least one group was performed followed by imputation with a downshift of 1.8 and a width of 0.3 standard deviations. Annotations were retrieved from UniProt for protein and gene names, Gene Ontology, and Keyword information. The data was visualized using principal component analysis (PCA) and proteins with significant changes in abundance between the respective comparisons were identified using volcano plots (Student’s *t*-test, p-value < 0.05; FDR = 5%, S_0_ = 1). 1D annotation enrichment heat maps defined changes in abundance across defined protein categories by Student’s *t*-test, p-value < 0.05; FDR = 0.05, and score <-0.5, >0.5^56^. Proteins were sorted by Gene Ontology (GO) Biological Processes (GOBP) and GO Molecular Function (GOMF), as well as UniProt Keywords. Box plots were generated with GraphPad Prism v9 and a Student’s t-test p-value < 0.05 was used for statistical testing relative to the untreated control. STRING network interaction mapping was used to visualize the interactions of proteins identified in the study and their cellular pathways^57^.

### SDS-PAGE and Western blot

Undigested protein extract from 1, 4, 8, 11, 15, and endpoint dpi were separated by 10% SDS-PAGE and stained with Coomassie to confirm protein loading. A second SDS-PAGE was transferred to a polyvinylidene fluoride membrane with the Trans-Blot Turbo Transfer System (Bio-Rad), according to manufacturer’s instructions. Membranes were blocked at 4 °C overnight in 5% non-fat skim milk in TBS (50 mM Tris, 150 mM NaCl, pH 7.5) followed by 2x washing with TBST (1x TBS, 0.05% Tween 20). Blots were incubated with the haptoglobin polyclonal antibody (Thermo Fisher Scientific) at 1:2,000 dilution in 5% non-fat skim milk in TBS at 4 °C overnight. Membranes were washed with 2x with TBST and incubated with 1:10,000 goat anti-rabbit IgG-horse-radish peroxidase secondary antibody (Thermo Fisher Scientific) for 1 h at room temperature. Next, blots were washed 6x with TBST and with the Cytiva Amersham™ ECL Select™ Western Blotting Detection Reagent (Fisher Scientific). The experiment was performed in technical duplicate.

## Results

### Increased depth of blood proteome coverage reveals previously undetected features of cryptococcal infection

To evaluate the power of proteomics toward improved diagnostic strategies for cryptococcal infection, we performed quantitative proteomics profiling on murine blood collected over the duration of disease (Fig. 1A). We confirmed progression of infection through a murine weight loss with a significant reduction in weight of infected mice (p-value = 0.0139) (Fig. 1B) correlated with the survival curve (Fig. 1C). We detected 19,118 – 22,981 peptides across uninfected samples and 19,678 – 27,186 peptides across infected samples with a marked increase in peptide identifications at the experimental endpoint (Fig. 1D). Correspondingly, we detected 2,892 – 3072 proteins across uninfected samples and 2,957 – 3,269 proteins across infected samples, with the largest differential between infection states at the endpoint (Fig. 1E). Prior to valid value filtering, we detected and quantified 3,067 host proteins and 160 fungal proteins (Fig. 1F). These findings demonstrate a 28-fold increase in detection of cryptococcal proteins within murine blood from our previous study^10^, supporting the potential of new technologies for improved biological insights.

**Figure 1:**
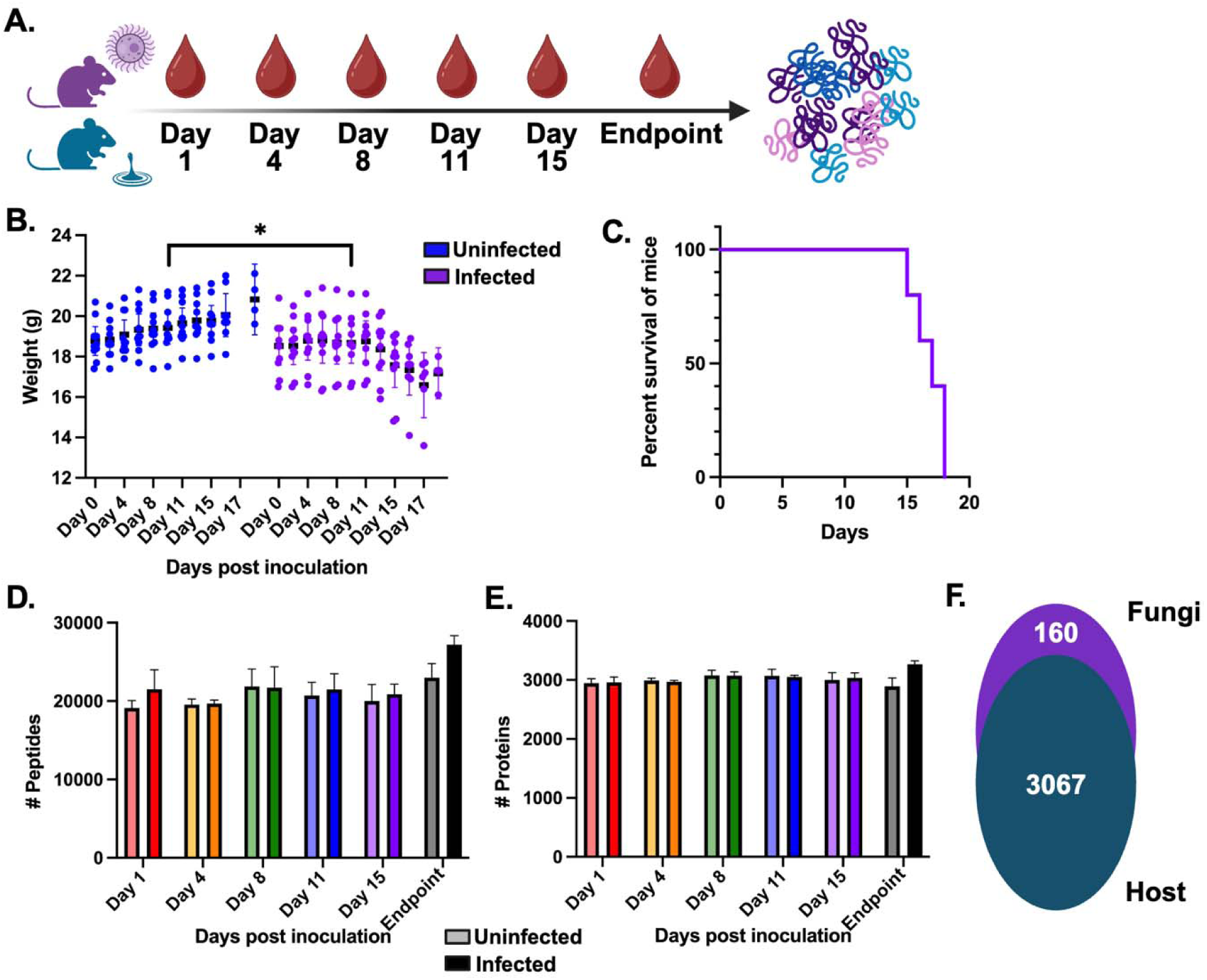
Experimental workflow and proteome overview. **A.** Murine infection model assay with *C. neoformans*-infection (purple) and no infection (blue). Blood collection at indicated time points followed by proteomics sample preparation and measurements. **B.** Weight distribution for each uninfected or infected mouse over the study duration (an uninfected control was matched to each endpoint collection), two-way ANOVA (mixed effects model) was applied, *p-value = 0.0139. **C**. Survival curve of *C. neoformans*-infected mice. **D.** Number of peptides measured (average +/- standard deviation) across five biological replicates per time point and condition. **E.** Number of proteins measured (average +/- standard deviation) across five biological replicates per time point and condition. **F.** Number of host (*M. musculus*) and fungal (*C. neoformans*) proteins identified and quantified after valid value filtering (i.e., presence of protein across 70% biological replicates within at least one group).

### Host blood proteome remodeling defines infection state and temporal drivers of disease response

Given the differences in protein detection and quantification levels reported above, we aimed to tease apart the impact of cryptococcal infection on the murine blood proteome over the duration of disease. Specifically, we prioritized the host core proteome (i.e., proteins detected in 70% of replicates within each group) for these insights. The principal component analysis (PCA) defines clustering by infection state (i.e., uninfected vs. infected samples) along component 1 accounting for 25.87% of data variance and the y-axis by time (i.e., 1, 4, 8, 11, 15, 18 dpi) for component 2 accounting for 10.07% of data variance (Fig. 2A). Next, we evaluated changes in abundance of individual host proteins at each time point between infected vs. uninfected samples (Fig. 2B). We identified one protein (alpha-2-HS-glycoprotein, AhsG) with increased abundance during infection compared to two proteins (murinoglobulin-2, Mug2; major urinary protein 18, Mup18) with increased abundance in the uninfected state at 1 dpi. The core host proteome did not show additional changes in protein abundance until 15 dpi with eight proteins significantly higher in abundance during infection compared to three proteins significantly lower. Critically, the proteins with elevated abundance during infection at 15 dpi captured known immune-associated proteins, including haptoglobin, eosinophil peroxidase, and immunoglobulins (Ighg1, Ighg, Ighg2b), whereas proteins with reduced abundance included apolipoprotein C-II, associated with maintaining lipid homeostasis^58^. At the experimental endpoint (i.e., 18 dpi), we detected 212 proteins with significantly increased abundance during infection compared to 95 proteins with significantly lower abundance (Supplemental Table 7).

**Figure 2:**
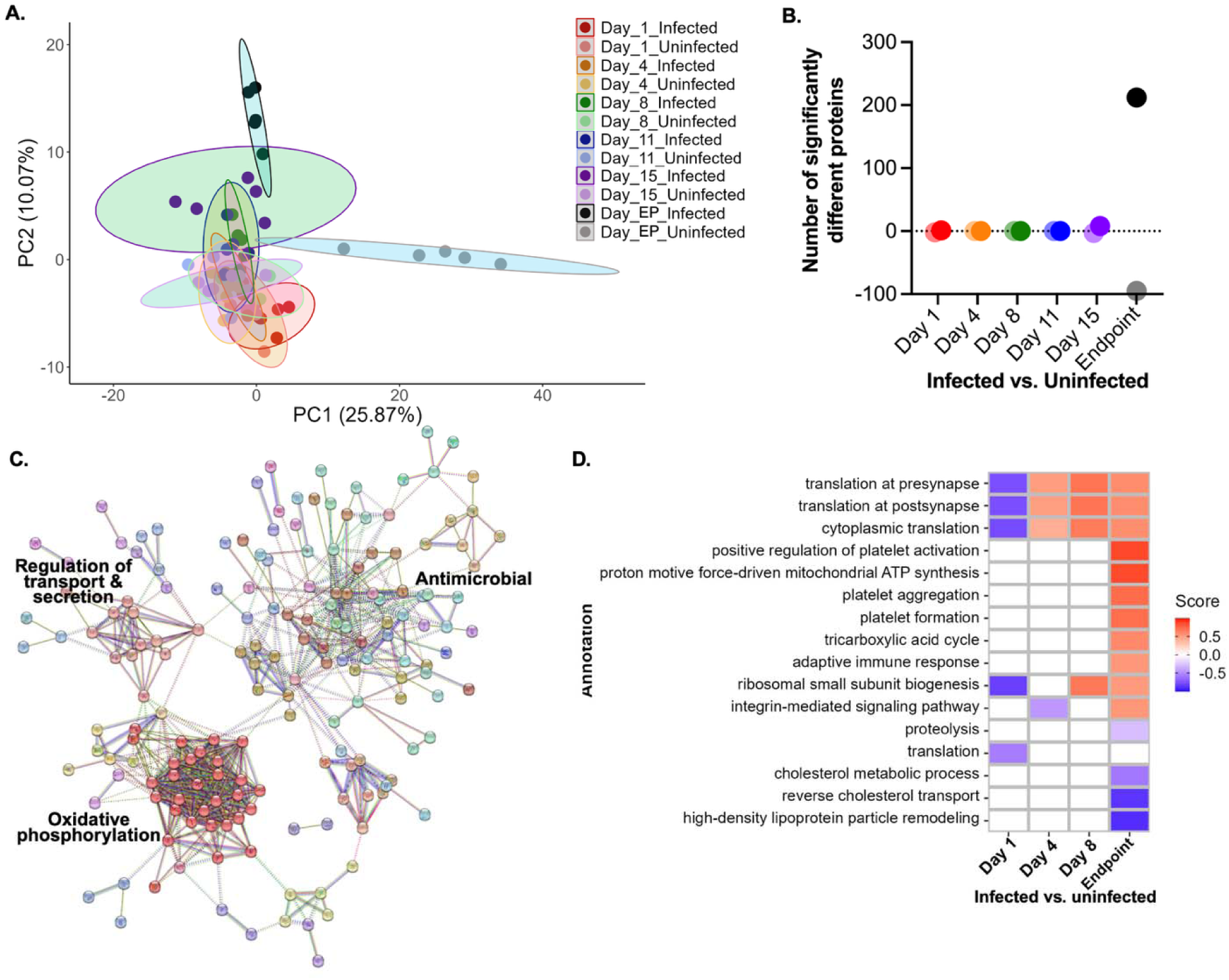
Temporal remodeling of the host core proteome. **A.** Principal component analysis. **B.** Volcano plot dot plot of significantly different proteins. Positive values = number of proteins significantly higher during infection. Negative values = number of proteins significantly higher during healthy (i.e., uninfected) state. **C.** STRING network diagram of host protein interactions for proteins with significant increase in abundance at endpoint during infection. MCL clustering (inflation parameter = 3; dotted link = edges between clusters; Evidence by text mining, experiments, databases, co-expression, neighbourhood, gene fusion, and co-occurrence; confidence [high] = 0.700; disconnected nodes in the network are removed). **D.** 1D annotation enrichment by Gene Ontology Biological Processes. All comparisons assessed by Student’s t-test p-value < 0.05; FDR = 5%; only comparisons with enrichment displayed.

Next, an investigation of the significantly elevated host proteins using STRING complemented with MCL (Markov Cluster Algorithm) clustering revealed functional roles by Gene Ontology of these proteins associated with antimicrobial properties, regulation of transport and secretion, and oxidative phosphorylation (Fig. 2C). Further assessment of global changes by 1D annotation enrichment, which evaluates if numerical values corresponding with selected annotation terms have a preference to be systematically larger or smaller than the global distribution of the values of all proteins^56^, revealed distinct enrichment at 1 dpi, 4 dpi, 8 dpi, and endpoint (Fig. 2D). Specifically, 1D annotation enrichment by Gene Ontology Biological Processes (GOBP) defined proteins associated with translation as higher in uninfected samples compared to infected samples at 1 dpi, with a reversal of this trend observed at the later timepoints, along with elevated adaptive immune response and integrin-mediated signaling pathway at the endpoint. Conversely, proteolysis, cholesterol metabolic process and transport, as well as lipoprotein particle remodeling were lowered at endpoint. Notably, platelet activation, aggregation, and formation were elevated at the endpoint, suggesting technical influence (i.e., blood coagulation upon collection) on protein abundance within these categories. Together, these data support remodeling of the core blood proteome across specific timepoints of cryptococcal infection, with both anticipated activation of immune-associated proteins, and identification of novel putative biomarkers distinguish early vs. late infections.

### Infection-specific host responses prioritize immune system activation and new signatures of cryptococcal disease

Moving beyond core host proteome response to cryptococcal infection, we focused on assessing unique drivers between infected vs. uninfected states across the temporal axis of disease. For this analysis, protein IDs were filtered by valid values detected in 70% of samples in at least one group. Within uninfected samples, we detected 2,260 proteins common to all time points and proteins exclusive to endpoint (Fig. 3A). Conversely, 2,503 proteins were common to all infected groups with no proteins exclusively present (Fig. 3B). Next, we performed a 1D annotation enrichment test for global comparisons of infected vs. uninfected samples and as anticipated, we observed an enrichment by Keywords for proteins associated with host immune response, including immunoglobulin, integrin, antiport, antimicrobial, respiratory chain, adaptive immunity, and ion transport (Fig. 3C). Within the uninfected samples, we observed an enrichment for proteins by Keywords associated with extracellular matrix, collagen, and serine esterase. A comparison of specific proteins with significantly different abundance between all infected vs. uninfected samples defined 11 proteins with increased abundance during infection and seven proteins with decreased abundance (Fig. 3D; Supplemental Table 8).

**Figure 3:**
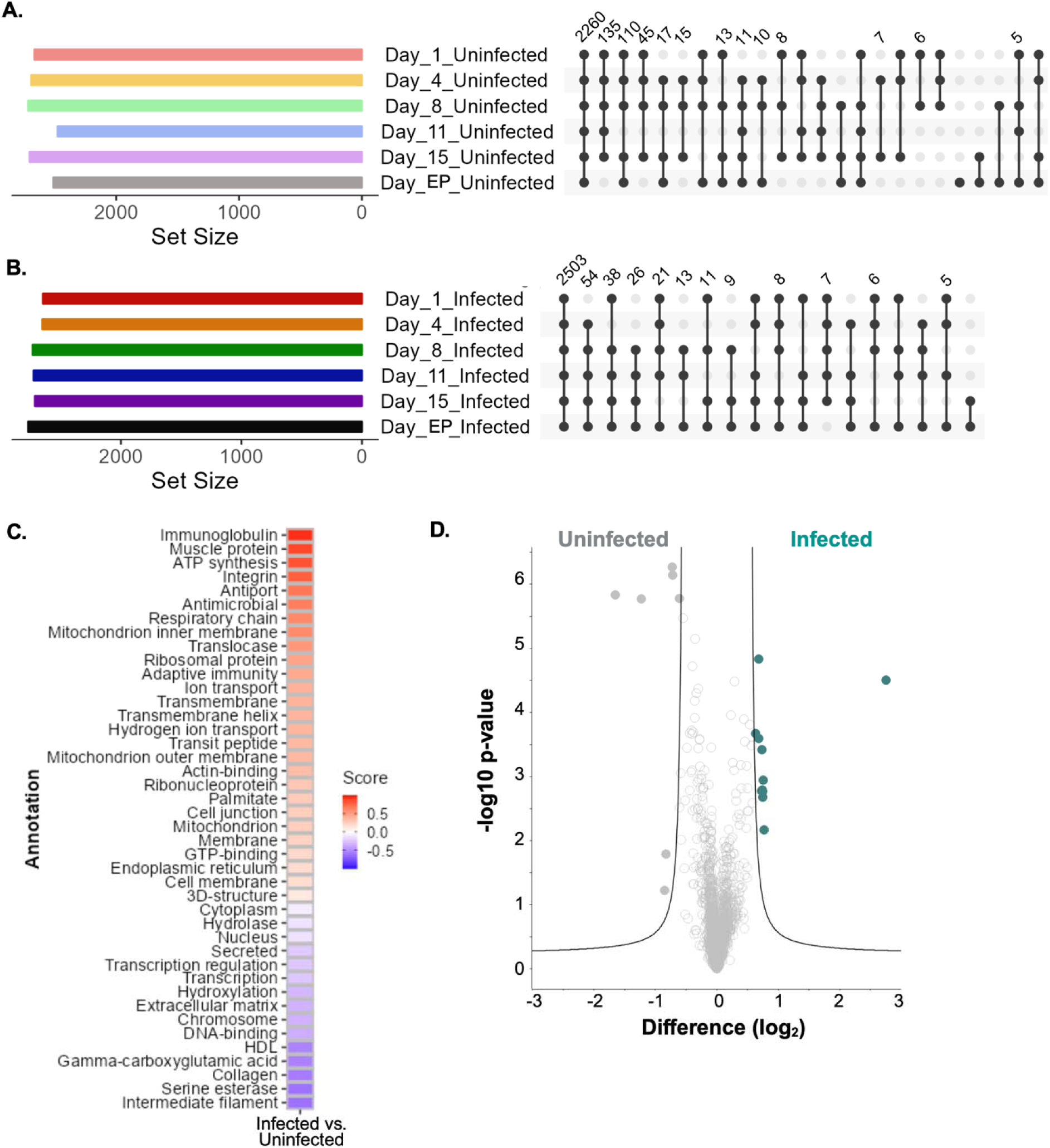
Infection-based host proteome remodeling. **A.** UpSet plot for host proteins identified across the uninfected samples. Note: repeat values are not indicated. **B.** UpSet plot for host proteins identified across the infected samples. Note: repeat values are not indicated. **C.** Heatmap of 1D annotation enrichment by Keywords (UniProt) for comparison of Infected (all time points) versus Uninfected (all time points). Student’s t-test p-value < 0.05; FDR = 5%. **D.** Volcano plot identifying significantly different proteins (Supplemental Table 8) between Infected (teal) and Uninfected (grey) samples across all time points. Student’s t-test p-value < 0.05; FDR = 5%; S_0_ = 1.

Assessing these significantly different proteins, we observed overlap with the temporal remodeling data and investigated abundance patterns at each time point. For instance, we identified two immunoglobulins (Igkv12, Ighg) with significantly elevated abundance during infection at 11 dpi, 15 dpi, and endpoint (Fig. 4A) and a single eosinophil peroxidase (Epo) with significantly increased abundance over time (excluding 4 dpi, where variance in the uninfected samples appears to influence significance) (Fig. 4B). Bone marrow proteoglycan 2 (Prg2; Fig. 4C), inter-alpha-trypsin inhibitor heavy chain H3 (Itih3; Fig. 4D), and three fibrinogen proteins (Fga, Fgb, Fgg; Fig. 4E) showed similar patterns of abundance with elevated levels during infection over time. With consideration of the later time points, we observed a significant increase in production of lipocalin-2 (neutrophil gelatinase-associated lipocalin, Lcn2; Fig. 4F), cathepsin G (CtsG; Fig. 4G), and haptoglobin (Hp; Fig. 4H). Importantly, given the role of haptoglobin as an acute phase protein involved in immune activation, oxidative homeostasis and hemoglobin scavenging, and pathogen opsonization^59,60^, we validated its increased production at endpoint via antibody-based detection (Fig. 4I) and provided semi-quantitation of haptoglobin abundance (Fig. 4J). Together, these data showcase the power of proteomics to identify host proteins with altered abundance profiles between infection states and across a time course of disease to propose novel biomarkers of cryptococcal infection from whole blood.

**Figure 4:**
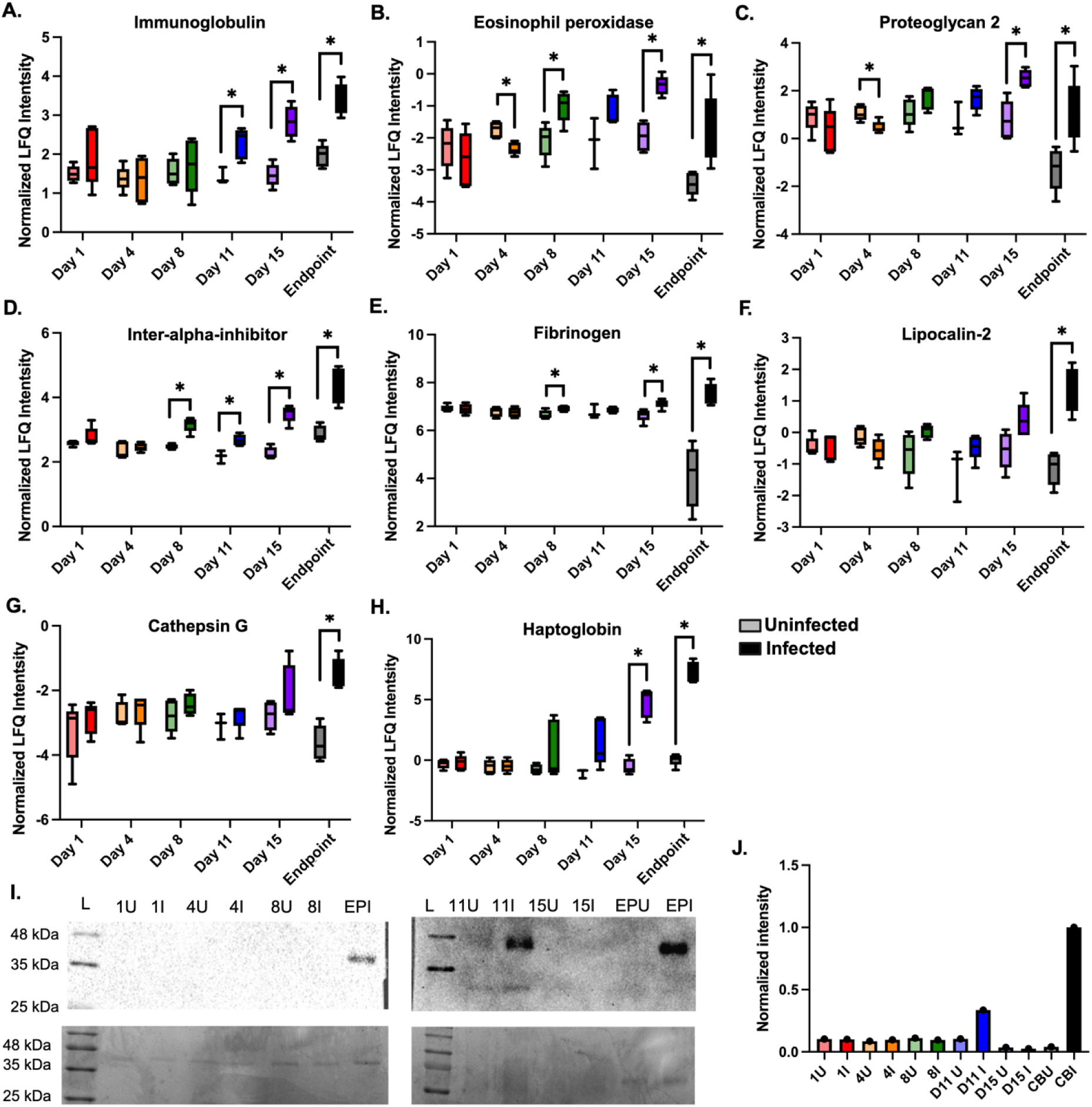
Immunomodulation within the host blood proteome upon cryptococcal infection. Boxplots of proteins with significantly higher abundance during infection compared to uninfected controls for **A.** Immunoglobulin (2 proteins). **B.** Eosinophil peroxidase**. C.** Proteoglycan 2. **D.** Inter-alpha inhibitor. **E.** Fibrinogen (3 proteins). **F.** Lipocalin-2. **G.** Cathepsin G. **H.** Haptoglobin. Statistical testing performed: Unpaired Student’s t-test, *p=value < 0.05. **I.** Western blot (top) of haptoglobin across uninfected (U) and infected (I) blood samples. Corresponding Coomassie-stained SDS-PAGE (bottom). **J.** Quantification of Western blot. Quantification performed within each blot and normalized to Endpoint I.

### In-depth fungal proteome mapping defines time-dependent remodeling and proposes traceable protein abundance patterns correlative with disease state

With improved detection of fungal proteins within a complex matrix (i.e., host blood), we can explore new mechanisms of host modulation and fungal pathogenesis. Based on valid value filtering of infected samples for 70% protein identification within at least one group, we defined a core fungal proteome (common within all groups) of 43 proteins and observed exclusive fungal protein production at 4 dpi and endpoint (Fig. 5A; Supplemental Table 9). Focusing on this core proteome, we observed a distribution of functional roles by GOBP for the fungal proteins associated with pathogenesis, including ion binding, protein folding, proteolysis, signaling, stress response, and transport (Fig. 5B). Next, we aimed to assess if these fungal proteins displayed changes in abundance across the time course of infection (Fig. 5C). Herein, we observed consistent abundance independent of time for 34 fungal proteins, including several proteins with known roles in fungal virulence, such as a non-specific serine/threonine protein kinase involved in thermotolerance^61^, adenylate cyclase involved in titan cell formation and capsule regulation^62,63^, peroxiredoxin involved in polysaccharide capsule production and reactive oxygen species reduction^64,65^, and transcriptional coregulator SSA1 involved in stress tolerance and melanin production^61,66^. In comparison, two proteins showed highly variable production over time, an ATP-dependent RNA helicase associated with *C. neoformans* extracellular vesicles^67^ and cell division control protein 42 with known roles in thermotolerance and mating^68,69^. We also identified four proteins with increasing protein abundance over time (i.e., importin, alpha- tubulin, DNA repair protein, and glyceraldehyde dehydrogenase) with no defined roles in fungal virulence, as well as three proteins with decreasing abundance over time (i.e., phosphoglycerate kinase, heat shock protein 60, chaperone DnaK) with the heat shock protein being regulated by Pka1 and elevated upon zinc replete growth conditions^70,71^ and the chaperone DnaK associated with iron homeostasis^72^. Lastly, we focused on fungal proteins with abundance patterns uniquely associated with temporal profiling, including an ATP synthase subunit previously detected in cerebral spinal fluid and extracellular vesicles^66,73^, which was only detected at 1 and 4 dpi (Fig. 5D). Whereas, in comparison, four proteins were detected only at 15 dpi and endpoint, including a cytoplasmic protein, phospholipid-transporting ATPase, serine/threonine-protein phosphatase, and peroxisomal ATPase with the kinase associated with interleukin (IL)-1β, IL-6, and IL-10 production^74^. Critically, these fungal proteome data provide clear indicators of infection independent of time (i.e., consistent production across the temporal axis) for general diagnostic power, along with traceable protein abundance patterns indicative of infection state (i.e., early vs. late) correlation with immune system activation to propose accurate, less invasive strategies to diagnosis and monitor cryptococcal disease.

**Figure 5:**
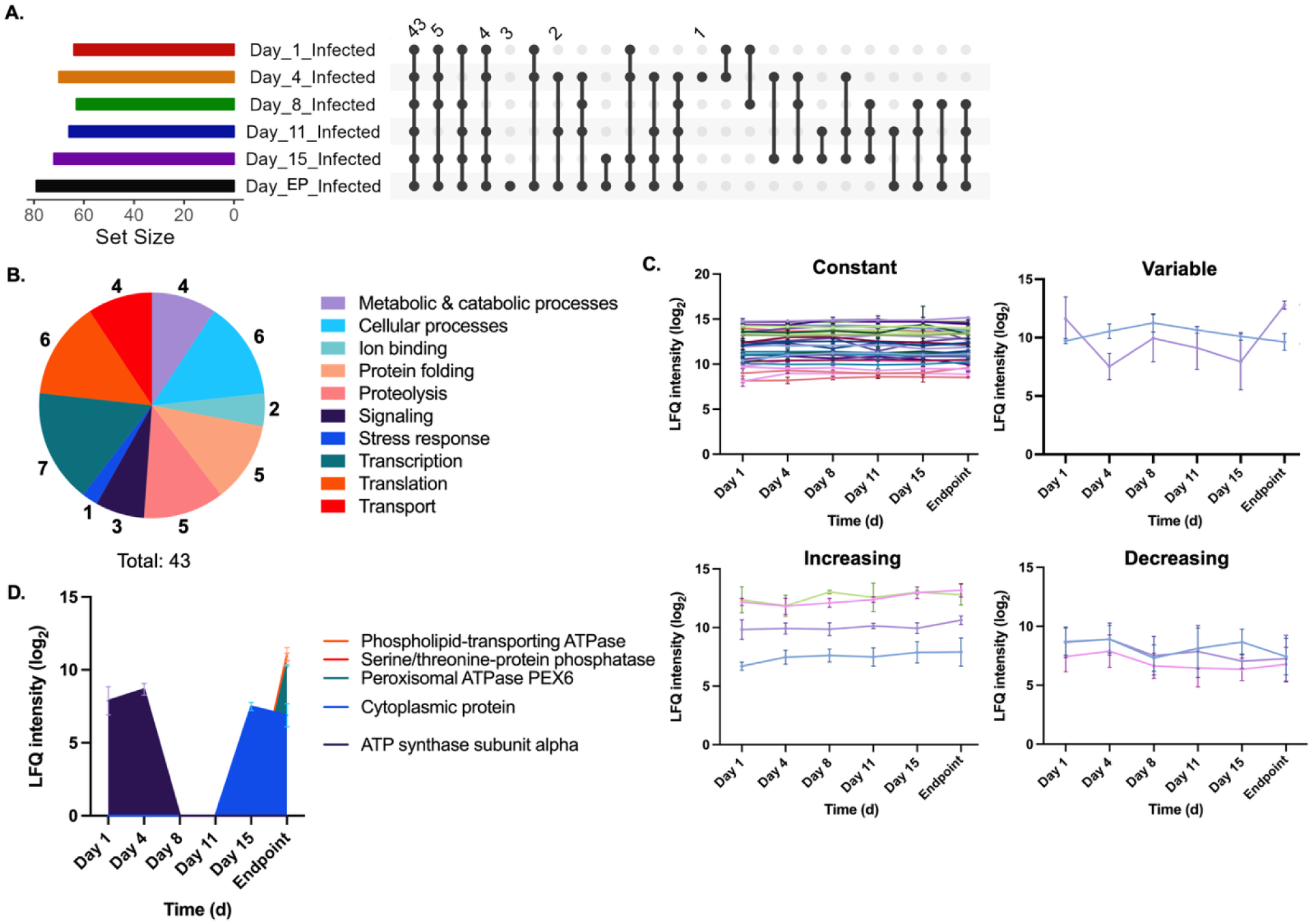
Fungal protein detection and modulation within the blood. **A.** UpSet plot for fungal proteins identified across the infected samples. Note: repeat values are not indicated. **B.** Pie chart of Gene Ontology Biological Processes terms for core fungal proteome (43 proteins). **C.** Abundance profiles of fungal proteins within the core proteome with specified production patterns across all time points: constant (i.e., unchanged abundance; 34 proteins), variable (i.e., abundance fluctuates over time; 2 proteins), increasing (i.e., abundance increases over time; 4 proteins), and decreasing (i.e., abundance decreases over time; 2 proteins). **D.** Abundance profiles of fungal proteins within the core proteome detected at early (i.e., days 1 and 4; 1 protein) and late (i.e., days 15 and endpoint; 4 proteins).

### Integration of mouse survival with host and pathogen protein abundance supports prognostic prediction of disease outcome

Based on our observations of activated host immune-associated proteins during cryptococcal challenge in a murine model across a time course of infection, novel modulation of haptoglobin by *C. neoformans*, and proposal of fungal virulence-associated signatures of disease, we aimed to assess the predictive power of dual perspective proteome profiling toward prognostics. We selected a mouse that survived until the end of the trial (mouse #6; 18 dpi) and a mouse that succumbed to infection at the earliest time point (mouse #7; 15 dpi) (Fig. 6A). For each comparison, we report the average value across all replicates. We correlated this difference in survival with weight loss between 0 dpi and endpoint showing -1.6% (mouse #6), -15.8% (mouse #7), and -5.1% (average) (Fig. 6B). Next, to evaluate a connection between host immune system activation and survival, we normalized protein abundance for infection-associated host proteins and observed a substantial increase in lipocalin-2, cathepsin G, and haptoglobin at 4 dpi for mouse #6 (Fig. 6C). Notably, mouse #7 showed elevated levels of immunoglobulins at 4 dpi but lower levels of all other infection-associated proteins. An assessment of immune system activation and sustainment at endpoint showed consistent elevation of lipocalin-2 and haptoglobin in mouse #6, along with immunoglobulins, and inter-alpha trypsin inhibitor (Fig. 6D). Conversely, mouse #7 showed an increase in eosinophil peroxidase production and cathepsin G at the endpoint but a substantial decrease in proteoglycan 2 and lipocalin-2.

**Figure 6:**
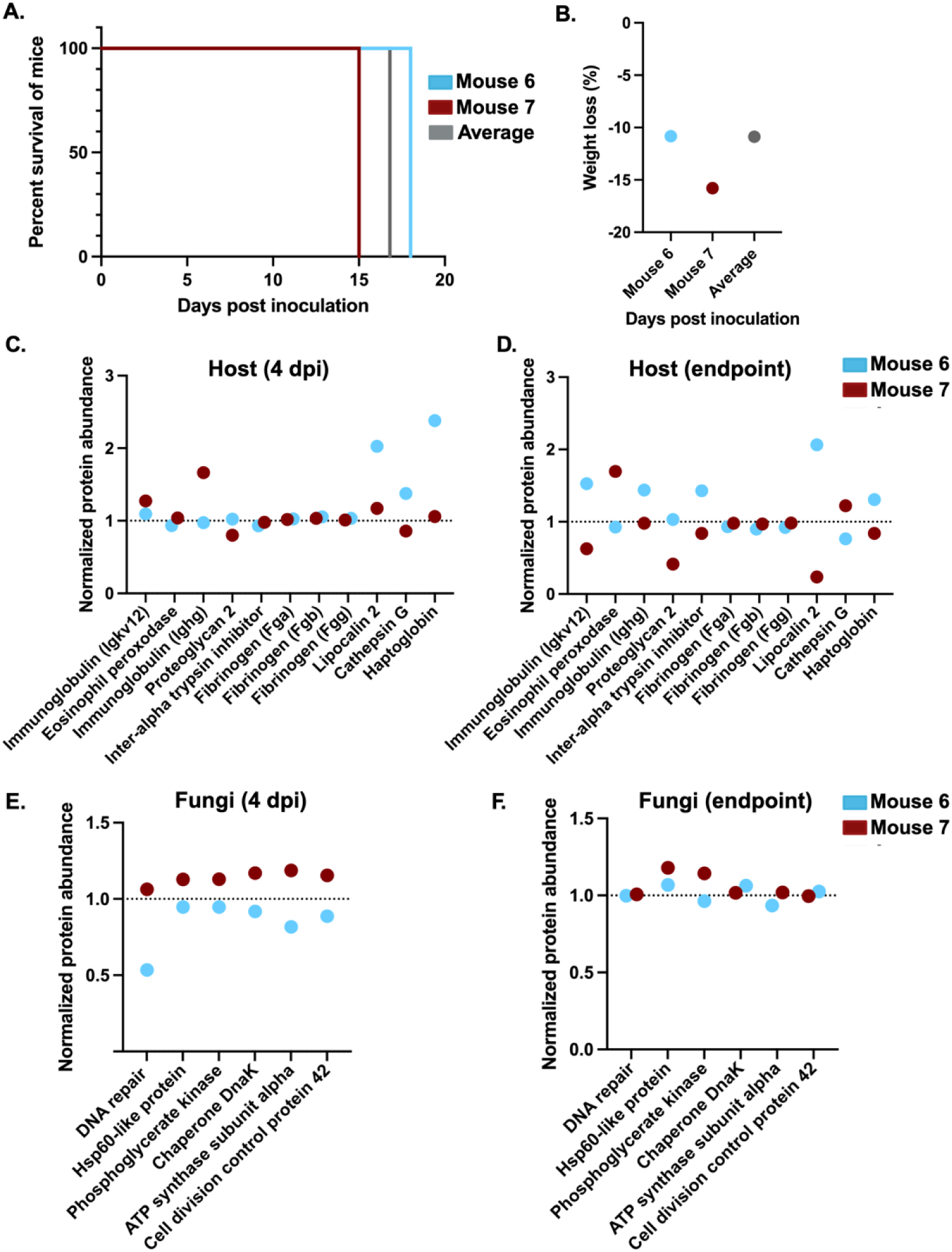
Integration of individual mouse survival data correlates with host immune response and fungal virulence. **A.** Survival curve for mouse #6 (survives until end of trial; 18 dpi), mouse #7 (reaches endpoint by 15 dpi), and average (survival rate for all infected mice in trial). **B.** Weight loss mouse #6 (-1.55%), mouse #7 (-15.79%), and average (-5.12%). Difference in weight loss considered between 0 dpi and endpoint. **C.** Normalized protein abundance plots for infection-associated host proteins at 4 dpi. Values normalized to the average value for each mouse. **D.** Normalized protein abundance plots for infection-associated host proteins at endpoint. Values normalized to the average value for each mouse. **E.** Normalized protein abundance plots for virulence-associated fungal proteins at 4 dpi. Values normalized to the average value for each mouse. **F.** Normalized protein abundance plots for virulence-associated fungal proteins at endpoint. Values normalized to the average value for each mouse. Average protein abundance normalized to 1.

Next, we assessed the impact of fungal protein production on individual mouse survival to observe consistently elevated abundance of virulence-associated proteins in mouse #7 at 4 dpi compared to a reduced abundance in mouse #6 (Fig. 6E). Similarly, at endpoint, mouse #7 still showed elevated abundance of HSP60-like protein and phosphoglycerate kinase compared to mouse #6 (Fig. 6F). Taken together, heightened levels of fungal virulence-associated proteins correlated with poor immune system activation beyond general immunoglobulins and correspond with reduced mouse survival. Conversely, elevated abundance of immune response proteins correlated with reduced fungal virulence to correspond with increased mouse survival. Overall, these data support the predictive prognostic power of dual perspective proteome profiling on individual survival.

## Discussion

The diagnostic and prognostic potential of quantitative proteomics is promising; however, its application towards infectious diseases, and specifically, fungal infections, is limited by detection of fungal proteins within the complex host matrix. Within this study, we use state-of-the-art mass spectrometry instrumentation to increase the depth of coverage and throughput for dual perspective complex proteome profiling from murine whole blood over a time course of cryptococcal infection. We define a core host proteome and identify immune-associated proteins with altered abundance over time, as well as host proteins with time-dependent abundance patterns. We validated detection of haptoglobin, an immune-associated protein involved in host oxidative homeostasis, opsonization, and hemoglobin scavenging upon host cell proteolysis, through Western blot in whole blood samples across time and infection state. Additionally, we correlated activation of haptoglobin by the host in response to proteolysis-associated proteins produced by *C. neoformans* throughout infection. Moreover, we propose putative biomarkers of cryptococcal infection based on protein abundance changes for both host and fungal proteins with several proteins displaying significantly altered profiles at early (i.e., 1 dpi) and late (i.e., endpoint) time points. Furthermore, we integrate knowledge of murine survival and weight loss (i.e., indicative of disease progression) with protein abundance profiles from the host and pathogen and demonstrate a correlative predictive power of heightened immune response and weakened fungal virulence. Together, these data demonstrate the predictive power of whole blood proteome profiling for individualized diagnostic and prognostic signatures of cryptococcal disease.

The detection of low abundant peptides and proteins from a contaminating pathogen within the background of highly abundant host proteins represents a major limitation of applying quantitative proteomics toward clinical research applications for infectious disease diagnostics^11,75^. In the absence of pathogen proteins, diagnostics rely on modulation of the host proteome, which is a promising approach in disease prediction and diagnosis for neurodegenerative disorders and cancer^76,77^, but limits the potential towards infectious disease as individuals may demonstrate unique baseline levels of pathogen exposure, depending upon environmental factors (e.g., urban vs. rural living) or predisposition for co-morbidities (e.g., immunocompromised status, co-infections). Importantly, technological developments in mass spectrometry, including the accessibility and adoptability of DIA measurement methods, library-free searching parameters in the absence of sequencing information or use of a data-dependent acquisition sample library, and instrumentation advances that increase sensitivity and speed of measurement, as well as limited ion loss, are pushing current boundaries^45,78–81^. Moreover, improvements to dynamic range measurements and quantification values (i.e., less missing values) support new discoveries across biological systems. Applications of quantitative proteomics toward fungal disease diagnostics are limited and therefore, within the current study, we performed proteome profiling using the Orbitrap™ Astral™ Zoom mass spectrometer. We detected over 3,000 host proteins from whole blood across a time course of cryptococcal infection and remarkably, we identified 160 fungal proteins, demonstrating an unprecedented depth of pathogen protein detection within the complex host matrix. Previous studies of fungal protein detection within whole blood focused on targeted proteomics measurement (i.e., multiple reaction monitoring) of specific peptides for the identification of three fungal proteins with diagnostic potential^10^. Given the global impact of fungal disease on human health, further investigation of the putative fungal protein signatures detected within the current study is warranted within clinical research samples to evaluate transition of the diagnostic potential from the lab to clinical research.

Across the host proteome, we uncover anticipated changes upon challenge with *C. neoformans* cells. For example, we observe the anticipated activation of the immune response, including elevated production of hypoxia-associated proteins, lipopolysaccharide binding protein, immunoglobulins, glycoproteins, integrins, and vesicle-trafficking proteins, elastases, lactotransferrin, myeloperoxidase, eosinophil peroxidase, and neutrophil granule proteins, S100- A8 and -A9, lipocalin, and haptoglobin, to highlight a few with defined roles in defense response, adaptive immune response, immune cell migration, and apoptosis^82–86^. For fungal pathogens, neutrophil elastases (e.g., Lipocalin-2) and myeloperoxidases regulate the formation of neutrophil extracellular traps (NETs), which are released from activated neutrophils and lead to microbicidal activity through the action of these granule proteins^87,88^. Specifically, NETosis pathways are stimulated by components of fungal pathogens, including the polysaccharide capsule and subsequent recognition of ß-glucans and mannans^89,90^. Lactotransferrin is an iron-binding glycoprotein that can limit fungal growth via a fungistatic effect through nutrient and iron sequestration within a limited environment, disrupt fungal cell surface permeability, and enhance antifungal drug efficacy^91,92^. We also observed a significant increase in S100A8 and S100A9, as well as lipocalin-2 at the experimental endpoint with each of these proteins previously correlated with protective anti-cryptococcal immune responses^93^. Specifically, for lipocalin-2, this serves as a potential biomarker derived from cerebral spinal fluid for bacterial meningitis^94^ and a differentiating component between bacterial and fungal meningitis^95^. Finally, the direct connection between cryptococcal infection and haptoglobin induction observed within this study is a novel finding that correlates free iron and heme availability and scavenging through the recycling of hemoglobin, as well as macrophage activation upon infection^96–99^. Herein, we show significant induction of haptoglobin within whole blood upon cryptococcal infection and propose its use as a diagnostic marker of disease.

From the pathogen perspective, we observe changes in *C. neoformans* protein abundance depending upon time and subsequent exposure to the host immune response with many proteins showing constant levels of production throughout infection, whereas the abundance and production of some proteins is directly correlated with infection state. For example, our previous characterization of HSP60-like protein correlated protein abundance with regulation by the cAMP/Protein Kinase A signaling pathway with primary roles in modulation of *C. neoformans* virulence, as well as increased abundance upon growth in zinc-limited media, supporting a role in nutrient acquisition and virulence^70,100,101^. Similarly, we identified an ATP synthase subunit alpha that was previously detected in the supernatant of *C. neoformans* culture media also under regulation of the cAMP/Pka pathway^10^. Within the current study, this protein was detected only during the early stages of infection, suggesting a role in fungal virulence and host modulation. Notably, we also identified several fungal proteins previously localized to the extracellular vesicles of *C. neoformans*, including the HSP60-like protein, ATP synthase subunit alpha, phosphoglycerate kinase, peroxiredoxin, transcriptional regulator SSA1, and glyceraldehyde-3-phosphate dehydrogenase, suggesting the release of vesicles during infection and subsequent dissemination throughout the blood^67,102^.

Finally, we propose a predictive model for the prognosis of cryptococcal infection by monitoring the abundance of host immune-associated and fungal virulence-associated proteins where an inverse relationship influences individual mouse survival. The application of proteomics toward disease prognostics has been explored in cancer but is limited within infectious disease research^103,104^. Specifically, in mouse #7, which succumbs to infection at the earliest time point (15 dpi), fungal virulence-associated proteins, such as HSP60-like protein, chaperone DnaK, and ATP synthase, show heightened abundance compared to the average profile and this correlates with a limited host immune response where only immunoglobulin activation is observed, along with low levels of lipocalin-2. These data propose a model of heightened fungal virulence in the absence of specific immune responses (i.e., haptoglobin, lipocalin-2, cathepsin), and the activation of immunoglobulins is insufficient to impact fungal virulence. These appear as critical factors driving survival of an individual mouse and propose a strategy to determine prognosis of infection based on these protein abundance patterns. Corroboratively, we observe the opposing pattern in mouse #6 where elevated immune response at early and endpoint correspond with reduced fungal virulence and increased murine survival. This information is assessed on an individual basis with the intent to highlight the potential of personal protein profiles to determine the prognosis of cryptococcal infection, a new parameter in disease management. Future studies will evaluate the disruption of a key immune-associated proteins (e.g., haptoglobin) on fungal virulence and subsequent mouse survival, as well as potential assessment within human clinical research samples evaluating the prognostic power of these dual perspective proteomes toward cryptococcosis.

## Conclusion

Herein, we perform the deepest dual perspective proteome mapping of cryptococcal infection from murine whole blood for the discovery of novel diagnostics signatures of host immune response and fungal virulence. We also highlight a new mechanism of immune system activation through production of haptoglobin in response to *C. neoformans.* Further, we demonstrate the predictive power of quantitative proteomics towards individual host survival through integration of immune system activation correlated to virulence suppression. Overall, we propose novel strategies for accurate and reproducible detection of cryptococcal infection from diagnostic biomarkers collected upon minimal invasion and we redefine disease management parameters through prognostic signature detection.

## Acknowledgements

The authors thank members of the Geddes-McAlister lab and Thermo Fisher Scientific for helpful discussions and constructive comments on the study.

## Author Contributions

M.W., J.A.M, J.D., S.N.S., & J.G.-M. conceptualized the study. M.W., J.A.M, J.D., A.H., D.H., J.R., S.N.S., & J.G.-M. designed the study. M.W., L.S., M.S., performed the murine infection assays and collected samples. M.W., J.A.M., & J.D. prepared and measured samples on the mass spectrometer. L.S. performed validation experiments. M.W., J.D., & J.G.-M. performed data analysis. J.D. contributed to manuscript methods and table preparation. J.G.-M. wrote the manuscript, generated figures and tables. All authors contributed to manuscript preparation and have read and approved the submitted manuscript.

## Data Availability

The proteomics datasets will be publicly available through PRIDE Proteomics Exchange. This submission includes a partial dataset. Additional RAW files will be added shortly after manuscript publication to complete the full data submission Project Accession: PXD064393 Token: 1l9utQ44QHYG Alternatively,

**Password:** KrFE0hI7GIRL

## Supplemental Files

**Supplemental Table 1. Liquid chromatography gradient and configuration.**

**Supplemental Table 2. Orbitrap Astral Zoom mass spectrometer global source and mass spectrometer parameters.**

**Supplemental Table 3. Orbitrap Astral Zoom mass spectrometer MS1 full scan experiment parameters.**

**Supplemental Table 4. Orbitrap Astral Zoom mass spectrometer MS2 DIA scan experiment parameters.**

**Supplemental Table 5. Orbitrap Astral Zoom mass spectrometer gas phase fractionation MS1 full scan experiment parameters.**

**Supplemental Table 6. Orbitrap Astral Zoom mass spectrometer gas phase fractionation MS2 DIA scan experiment parameters.**

**Supplemental Table 7. Significantly different core host proteins upon comparison to corresponding uninfected samples across the infection time course.**

**Supplemental Table 8. Significantly different host proteins during comparison of combined infected vs. uninfected samples.**

**Supplemental Table 9. Fungal proteins identified across infected samples following valid value filtering (>70% in at least one group).**

