## Supplemental Table 1-6 for "Whole blood proteome dynamics defines predictive diagnostic and prognostic signatures of cryptococcal infection"

**Supplemental Table 1: Liquid chromatography gradient and configuration.**

| 60 SPD Gradient Specifications and LC Configuration | | | |
| --- | --- | --- | --- |
| Gradient | Time (min) | % Mobile Phase B | Flow (μl/min) |
|  | 0 | 10 | 2.0 |
|  | 0.3 | 10 | 2.0 |
|  | 0.6 | 10 | 0.8 |
|  | 13.6 | 22.5 | 0.8 |
|  | 20.5 | 35.0 | 0.8 |
|  | 20.9 | 55.0 | 2.0 |
|  | 20.95 | 99.0 | 2.0 |
|  | 22.35 | 99.0 | 2.0 |
| LC Parameters | LC Configuration | Trap and Elute | |
|  | Fast Loading/Equilibration Mode | Pressure Control | |
|  | Loading/Equilibration/Wash Pressure | Max Pressure | |
|  | Equilibration Factor | 3 | |
|  | Sampler Temperature | 7 °C | |
|  | Mobile Phase A / Weak Wash | 0.1% Formic Acid in Water | |
|  | Mobile Phase B / Strong Wash | 0.1% Formic Acid in 80% Acetonitrile | |
|  | Zebra Wash | Enabled | |
|  | Zebra Wash Cycles | 4 | |
|  | Analytical Column Temperature | 50 °C | |
| Column Specifications | Analytical Column | EASY-Spray™ PepMap™ Column, 2µm C18, 150µm × 15 cm (P/N ES906) | |
|  | Trap Column | PepMap™ Neo Trap Cartridge, 5 μm C18 300 μm x 5 mm, (P/N 174500) | |

**Supplemental Table 2. Orbitrap Astral Zoom mass spectrometer global source and mass spectrometer parameters.**

| Global Parameters (Source & MS) | |
| --- | --- |
| Positive Ion Voltage | 2100 Volts |
| Ion Transfer Tube Temperature | 290 °C |
| Expected Peak Width | 10 seconds |
| Default Charge State | 2 |
| Lock Mass Correction | Off |

**Supplemental Table 3. Orbitrap Astral Zoom mass spectrometer MS1 full scan experiment parameters.**

| MS1 Full Scan Experiment Parameters | |
| --- | --- |
| Orbitrap Resolution | 240K |
| Scan Range (*m/z*) | 380-980 |
| RF Lens (%) | 40 |
| Normalized AGC Target (%) / Absolute AGC Value | 500% / 5.00e6 |
| Maximum Injection Time | 3 milliseconds |
| Microscans | 1 |

**Supplemental Table 4. Orbitrap Astral Zoom mass spectrometer MS2 DIA scan experiment parameters.**

| MS2 DIA Scan Experiment Parameters | |
| --- | --- |
| Precursor Mass Range (*m/z*) | 380-980 |
| Isolation Window (*m/z*) | 3 |
| Window Placement Optimization | On |
| AGC Target | Custom |
| Normalized AGC Target (%) / Absolute AGC Value | 500% / 5.00e4 |
| Maximum Injection Time | 7 milliseconds |
| DIA Scan Range (*m/z*) | 150-2000 |
| HCD Collision Energy (%) | 25 |
| RF Lens (%) | 40 |
| Pre-Accumulation | On |
| Loop Control | Time |
| Time | 0.6 seconds |

**Supplemental Table 5. Orbitrap Astral Zoom mass spectrometer gas phase fractionation MS1 full scan experiment parameters.**

| GPF MS1 Full Scan Experiment Parameters | |
| --- | --- |
| Orbitrap Resolution | 240K |
| Scan Range (*m/z*) | Incremental 100 *m/z* scans (380-480; 480-580; 580-680; 680-780; 780-880; 880-980) |
| RF Lens (%) | 40 |
| Normalized AGC Target (%) / Absolute AGC Value | 500% / 5.00e6 |
| Maximum Injection Time | 3 milliseconds |
| Microscans | 1 |

**Supplemental Table 6. Orbitrap Astral Zoom mass spectrometer gas phase fractionation MS2 DIA scan experiment parameters.**

| GPF MS2 DIA Scan Experiment Parameters | |
| --- | --- |
| Precursor Mass Range (*m/z*) | Incremental 100 *m/z* scans (380-480; 480-580; 580-680; 680-780; 780-880; 880-980) |
| Isolation Window (*m/z*) | 1 |
| Window Placement Optimization | On |
| AGC Target | Custom |
| Normalized AGC Target (%) / Absolute AGC Value | 500% / 5.00e4 |
| Maximum Injection Time | 18 milliseconds |
| DIA Scan Range (*m/z*) | 150-2000 |
| HCD Collision Energy (%) | 25 |
| RF Lens (%) | 40 |
| Pre-Accumulation | On |
| Loop Control | Time |
| Time | 0.6 seconds |
